# When the microbiome defines the host phenotype: selection on vertical transmission in varying environments

**DOI:** 10.1101/2020.09.02.280040

**Authors:** Marjolein Bruijning, Lucas P. Henry, Simon K.G. Forsberg, C. Jessica E. Metcalf, Julien F. Ayroles

**Affiliations:** Princeton University, Department of Ecology and Evolutionary Biology, 08544 Princeton, New Jersey, United States of America; Lewis-Sigler Institute for Integrative Genomics, 08544 Princeton, New Jersey, United States of America

**Keywords:** Bet hedging, fitness, fluctuating selection, holobiont, microbiome heritability, quantitative genetics

## Abstract

The microbiome can contribute to variation in fitness-related traits of their hosts, and thus to host evolution. Hosts are therefore expected to be under selection to control their microbiome, for instance through controlling microbe transmission from parents to offspring. Current models have mostly focused on microbes that either increase or decrease fitness. In that case, host-level selection is relatively straightforward, favouring either complete or no inheritance. In natural systems, however, vertical transmission fidelity varies widely, and microbiome composition is often shaped by a combination of vertical and horizontal transmission modes. We propose that such mixed transmission could optimize host fitness under fluctuating environments. Using a general model, we illustrate that decreasing vertical transmission fidelity increases the amount of microbiome variation, and thus potentially phenotypic variation, across hosts. Whether or not this is advantageous depends on environmental conditions, how much the microbiome changes during host development, and the contribution of other factors to trait variation. We discuss how environmentally-dependent microbial effects can favor intermediate transmission, review examples from natural systems, and suggest research avenues to empirically test our predictions. Overall, we show that imperfect transmission may be adaptive by allowing individuals to ensure phenotypic variability in their offspring in contexts where varying environments mean that this strategy increases long-term fitness.

## Introduction

Microbial life occupies almost every habitat on Earth. Increasingly, there is evidence that microbial communities living in and on eukaryotic hosts can strongly affect host phenotypes, shaping features including behavior (Bercik *et al.* 2011; Johnson & Foster 2018; Sherwin *et al.* 2019), development (Blanton *et al.* 2016; Charbonneau *et al.* 2016), illness (Matsuoka & Kanai 2015), pathogen resistance (Wei *et al.* 2015; Niu *et al.* 2017; Berg & Koskella 2018) and life span (Keebaugh *et al.* 2018). Such strong effects on host traits indicate that the microbiome can affect host fitness. Moreover, the composition of microbiome communities often varies greatly between hosts within a population, explaining a substantial proportion of host phenotypic variation (e.g. Camarinha-Silva et al., 2017; Difford et al., 2018). The importance of the microbiome for host fitness, together with the considerable variation between hosts, imply that, in theory, the microbiome has the potential to impact host adaptive evolution. However, as of yet, it is largely unknown how much the microbiome contributes to host adaptation (Moran & Sloan 2015; Henry *et al.* 2019).

The importance of the microbiome for host fitness implies that hosts will be under selection to ‘manage’ their microbiome communities (Foster *et al.* 2017): adaptations that enable hosts to control their microbiome composition have clear potential to increase fitness. Such adaptations could act on different stages during microbe acquisition and establishment (discussed by Foster, Schluter, Coyte, & Rakoff-Nahoum, 2017). Our focus here is on hosts controlling their microbe composition by controlling transmission of microbes from parents to offspring (i.e. vertical transmission). There exists a wide variety of mechanisms for transmission, producing a large diversity in the range of transmission fidelity across systems (Fig. 1). Some host species have faithful microbial transmission (Fig. 1A,B), leading to high concordance between the microbiomes of parents and offspring, ultimately echoing the inheritance of host genetic material. The most faithful transmission method is through intracellular infection of oocytes, epitomized in obligate nutritional symbiosis observed in many sap-feeding insects (Baumann 2005). For example, aphids are nearly all infected with *Buchnera* bacteria, enabling aphids to feed on phloem sap, an otherwise unbalanced diet (Douglas 1998). Other forms of vertical transmission occur through ‘intimate neighbourhood transmission’ during seed formation, egg laying, or passage through the birth canal (Roughgarden *et al.* 2018). Indeed, in order to transmit bacteria from parents to offspring, stinkbugs can attach special symbiont capsules to their eggs (Fukatsu & Hosokawa 2002) or cover eggs with symbiont-supplemented jelly (Kaiwa *et al.* 2014). Feces consumption (cophrophagy) is an important mechanism by which early-stage cockroaches acquire their gut bacteria, increasing their fitness compared to when reared under sterile conditions (Jahnes *et al.* 2019). In dung beetles, vertical transmission is ensured through a brood ball, which results in remarkably faithful microbial transmission (Estes *et al.* 2013). Recent modeling studies suggest that whenever transmission is faithful, as in these examples, the microbiome has the potential to contribute to host adaptation (Zeng *et al.* 2017; van Vliet & Doebeli 2019).

**Figure 1:**
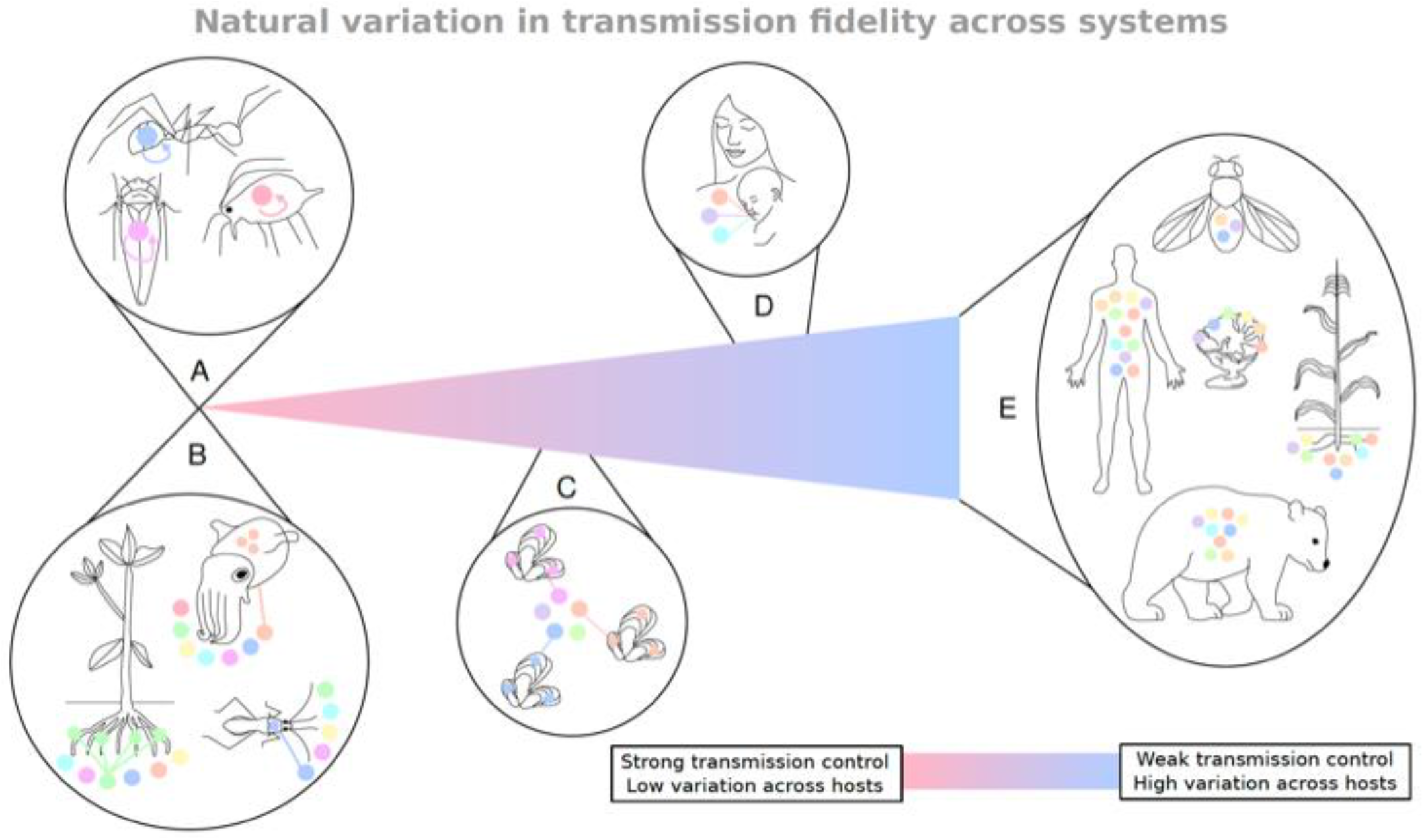
Natural variation in microbiome transmission fidelity across the eukaryotic kingdom, resulting in variation in microbiome composition across hosts in a population. A) Obligate intracellular symbionts represent strong control over microbial transmission. Across insects, such as carpenter ant-*Blochmannia* (Sauer et al. 2002), aphid-*Buchnera* (Koga et al. 2012), leafhopper-*Sulcia* associations (Brentassi et al. 2017), both host and microbe have intricate molecular mechanisms where symbionts are transovarially transmitted from mother to offspring, limiting symbiont diversity. B) Other hosts have similarly intricate mechanisms to acquire their microbes from the environment. Bobtail squid-*Vibrio* (Nyholm and McFall-Ngai 2004), bean bug-*Burkholderia* (Kikuchi et al. 2007), and legume-rhizobia (Ibez et al. 2017) systems display similar specificity to limit environmental acquisition to their particular microbial associations. While A and B demonstrate distinct mechanisms, with different evolutionary properties, they both result in low microbiome variation across hosts in a population. C) Hosts may exert strong control over which microbes may infect, but maintain flexible associations with broader diversity of microbes than in scenarios illustrated in A or B. For example, deep sea mussels restrict acquisition to single bacterial species, but this bacterial species may vary across individuals and populations (Picazo et al. 2019). Bacterial diversity is reduced within hosts, but variable across hosts. D) Some hosts have behavioral mechanisms that transmit only some portions of the microbiome. In humans, mother transmit a distinct subset of microbes to their infants that likely help with lactose digestion and immune development (Asnicar et al. 2017; Ferretti et al. 2018; Yassour et al. 2018). However, the homogenization of microbiota between mother and infant disappears over the next few years (Gilbert et al. 2018; Korpela et al. 2018). E) For many hosts, microbiome transmission is thought to be unfaithful, leading to high variance among individuals. In *Drosophila* and maize, only a small percentage of the microbiome is faithfully transmitted (Walters et al. 2018; Douglas 2019). Sponges harbor specific microbial communities, but are no more likely to share symbionts between parents and offspring as they are between species; rather sponge microbiomes are environment-specific (Bjork et al. 2019). For some hosts, like bears, flexibility in the microbiome may enable microbes associated with increased nutrient acquisition in preparation for hibernation (Sommer et al. 2016). This flexibility may also happen in humans (David et al. 2014), but drivers of microbial diversity in humans, and these other systems, is not well understood.

However, despite these intriguing examples of tightly linked host-microbe associations, most host populations have microbiomes that vary through time and across hosts, leading to hosts associating with variable microbial communities (Fig. 1C-E). For instance, vertical transmission in marine sponges is highly unfaithful; they are no more likely to share symbionts between parents and offspring as they are between species (Björk *et al.* 2019). Indeed, the heritability of bacteria genera seems to be highly variable, although most estimates come from domesticated systems: estimates range between 0.15-0.25 for root-associated microbes in maize (Walters *et al.* 2018); in cows and pigs, only 8 out of 144, and 8 out of 49 microbe genera, respectively, showed a heritability larger than zero (Camarinha-Silva *et al.* 2017; Difford *et al.* 2018). From these genera, heritability estimates ranged between 0.17-0.25 in cows (Difford *et al.* 2018) and between 0.32-0.57 in pigs (Camarinha-Silva *et al.* 2017). The generally low heritability has led some studies to suggest that selection at the level of the host and microbiome together, is unlikely to drive adaptive changes in most natural host systems (Moran & Sloan 2015; Douglas & Werren 2016).

While low heritability could indeed reflect a relatively small role of the microbiome for host adaptation, it could also be that current models lack relevant elements. One of these elements might be environmental variation. There is some empirical evidence suggesting that loose host-microbiome associations, implying low heritability, might be beneficial under changing environmental conditions. In animals that undergo metamorphosis, like holometabolous insects, flexibility in the microbiome may optimize phenotypes for different environments at different life stages (Hammer & Moran 2019). Further, the microbiome potentially plays a role in the variance in timing of important life history events, which could help hosts hedge their bets in unpredictable environments or when in competition (Metcalf *et al.* 2019). Finally, a study on wild marine sponges suggested that the observed unfaithful vertical transmission rates might benefit larvae facing variable environments (Björk *et al.* 2019). However, as of yet, such fluctuating selection has not been incorporated in models of how faithful microbial transmission should be to optimize host fitness.

In this perspective, we explore how vertical microbiome transmission fidelity could affect long-term host fitness, and how this interacts with constant versus fluctuating selection. Can the microbiome contribute to host adaptation, even if heritability is low, or perhaps *because* heritability is low? To tackle this question, we model host phenotypes as a function of their microbiome, and map host phenotypes to fitness, in interaction with the environment. We vary vertical transmission fidelity, and assess how this fidelity affects phenotypic distributions and long-term fitness of a population of hosts. Results reveal that decreasing vertical transmission fidelity increases the amount of microbiome variation, and thus phenotypic variation, among hosts (Fig. 2). We discuss how this is conceptually similar to genetic mutations, where higher mutation rates (analogous to lower microbiome transmission fidelity) increase variation among individuals. We lay out how increased variation among hosts can be either advantageous or disadvantageous, depending on environmental fluctuations and predictability. Increasing host phenotypic variation by imperfect transmission reflects a diversified bet hedging strategy (“don’t put all your eggs in one basket”): although increased phenotypic variation among genetically related individuals decreases the expected year-to-year individual fitness, a reduction in fitness variance across generations, may optimize a genotype’s fitness over the long-term (Philippi & Seger 1989) (Fig. 2). We then explore how selection on transmission fidelity is additionally shaped by the contribution of other factors to phenotypic host variation, and by the degree to which the microbiome changes during host development. Our study shows that the microbiome has the potential to contribute to host adaptation, not only by altering the mean phenotype to one that maps to higher fitness, but also potentially by adjusting the variance in phenotypes to increase fitness (Bull 1987; Bruijning *et al.* 2020). Under sufficiently large fluctuating selection, a low microbiome transmission fidelity, increasing host phenotypic variation, can benefit host fitness. Our findings provide a new lens to interpret a fast growing body of literature suggesting that the microbiome has the potential to contribute to host adaptation.

**Figure 2:**
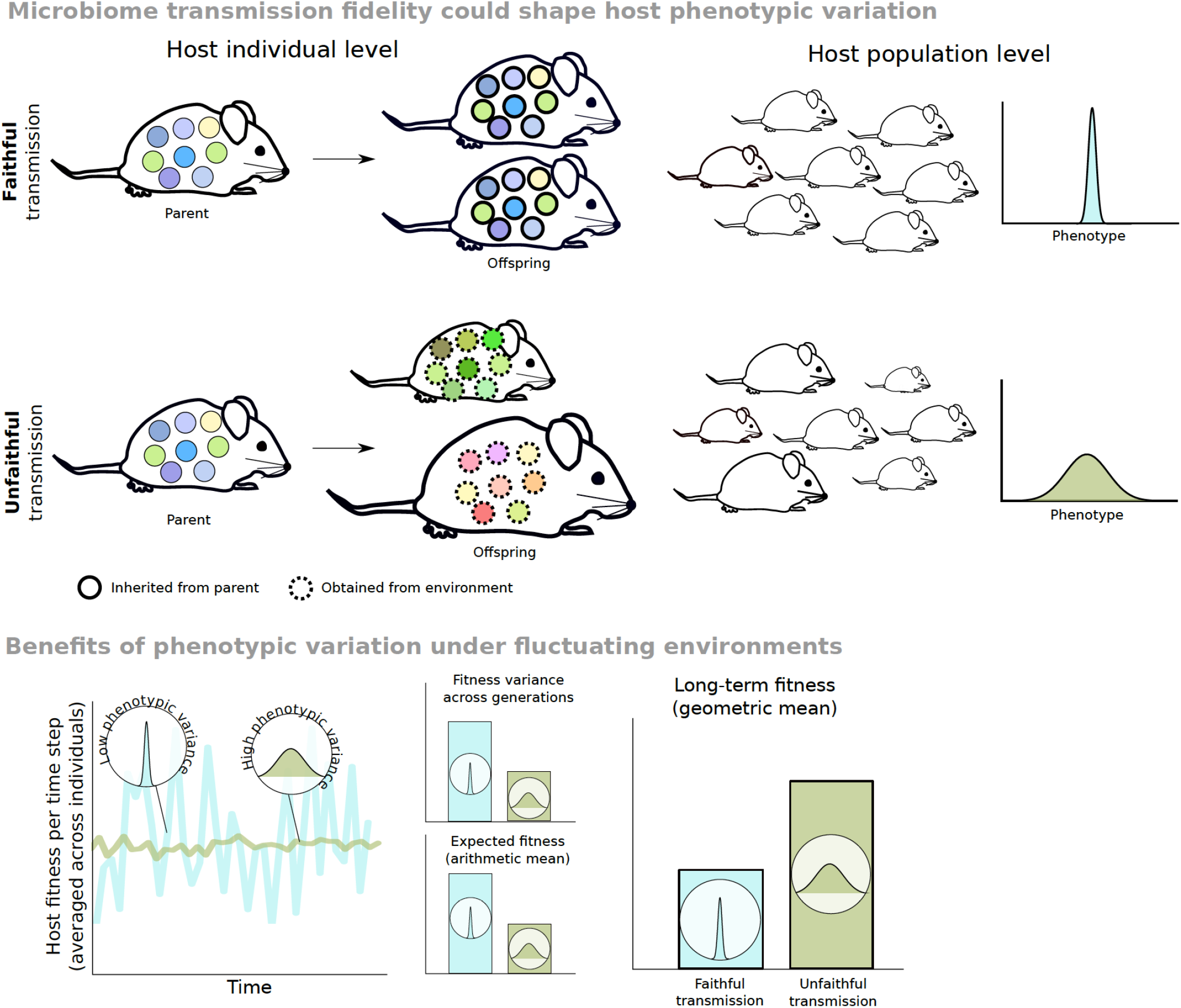
Conceptual overview of how microbiome transmission could shape host phenotypic variance, and when phenotypic variation among genetically similar individuals might be beneficial. Hosts are represented as mice and are characterized by their microbiome (circles). Microbes differ in their host effects (colors), and their combined effects determines their host’s phenotype (body size in this case). A population of hosts with faithful microbiome transmission will eventually result in the loss of phenotypic variation, due to stochasticity and selection. In a host population with unfaithful transmission, new variation is introduced every generation, resulting in maximal phenotypic variation. Increased phenotypic variation by imperfect transmission reflects a diversified bet hedging strategy. Under fluctuating environments, a reduction in fitness variance across generations, may optimize long-term fitness, despite a reduction in the expected year-to-year individual fitness. Host genotypes with low transmission fidelity could therefore be favored by natural selection.

## Modeling framework

### General overview

We use a modelling approach to assess how different factors shape host selection on vertical microbe transmission fidelity (Fig. S1). To do so, we run simulations for a range of parameters, reflecting different biological scenarios. In each scenario, we simulate individual hosts in a population, where a host’s microbiome composition affects its phenotype, which then shapes reproduction, upon which the next host generation takes place (Fig. S1A). These relations are governed and modified by different processes that vary across scenarios, such as the transmission fidelity, environmental conditions and microbial dynamics (see colored boxes in Fig. S1A; explained in detail in the next sections). For each simulation, we keep track of population-level outcomes: phenotypic distributions (both mean and variance) and fitness (Fig. S1B), averaged across time steps. An interactive tool to run the model can be found at http://marjoleinbruijning.shinyapps.io/simulhostmicrobiome, and R-code is available at http://github.com/marjoleinbruijning/microbiomeTransmission. Our model is conceptually and mathematically similar to quantitative genetic mutation-drift-selection models (see also Ravel, Michalakis, & Charmet [1997] for the case of a single symbiont), which describe how the balance between drift, selection and new mutations determines the amount of additive genetic variance in a population (Appendix S2).

In short, the simulation procedure is as follows. We simulate a microbial ‘source pool’, assumed to be present in the host environment, consisting of 1000 different microbial species, where each microbial species is equally abundant. Microbial species are characterized by their (additive) effects on host phenotypes. The effect that microbial species *j* has on its host phenotype is drawn from a normal distribution:

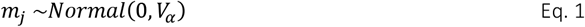

Here, *V*_*α*_ is the variance in microbial effects on host phenotype. We start by assuming that hosts do not differ in any other aspect. Thus, phenotypic variation among hosts results from variation in microbiome composition only. We simulate the dynamics of *N* hosts, where each host carries 100 microbes. At the start of a simulation, we assign 100 randomly sampled (with replacement) microbes from the source pool to each host. Throughout the simulations, both the number of microbes within each host, and the number of hosts, are kept constant, so that responses are due to changes in composition (of both hosts and microbes), and not due to changes in numbers. We follow dynamics across *T*_*h*_ host generations. Within each host generation, *T*_*m*_ microbial generations take place to capture changes in microbiome composition during host development (see section ‘*Within-host microbial dynamics*’). To explore the impact of temporal variability in the environment, every host generation, the relationship between phenotypes and fitness (see section ‘*Host selection and transmission of microbes*’) can change. This is captured by drawing from a normal distribution defined by:

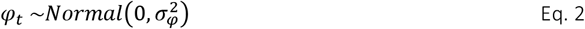

where 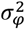 controls the degree of environmental fluctuations (when set at 0, the environment is stable). Eq. 2 simulates environments without any autocorrelation (*ρ*=0), i.e. the environment at time *t* is not predictive for the environment at time *t+1*. Below, we extend this by simulating environments using an autoregressive (AR) model with specified autocorrelation.

### Within-host microbial dynamics

To simulate microbial dynamics within a host generation, we use a metacommunity model. Each microbial generation, all microbes are replaced by either colonization (by immigration from the source pool outside the host) with probability c, or proliferation (replication of a microbe within the host microbe community) with probability 1-c (green box in Fig. S1A). We assume that all microbe species have equal colonization and proliferation probabilities, and that each microbial species is equally represented in the source pool. Such neutral dynamics have been shown to adequately describe microbial community composition in different systems (Sloan et al. 2006; Burns et al. 2016; Sieber et al. 2019). We note, however, that the model could be easily extended to allow for variation in these rates to model different microbe strategies, such as tradeoffs between colonization and proliferation rates, pathogenic strains with high proliferation rates, or variation in microbial abundance. For each host, this random colonization and proliferation is repeated for *T*_*m*_ time steps, allowing for changes in microbiome composition during host development. All hosts survive during this period.

### Host selection and transmission of microbes

After *T*_*m*_ time steps, host selection takes place based on host phenotypes, in interaction with the time-specific environment and the strength of selection (red box in Fig. S1A). The phenotype *P* of host *i* is calculated as the sum of all the effects of microbes that are present at the moment of host selection:

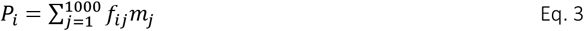

*f*_*ij*_ is the number of times microbe *j* is present in host *i* at the moment of selection (note that summing *f*_*ij*_ over all *j* equals 100; the total number of microbes within each host). This procedure implies that microbial composition earlier in life has no effect on current host phenotype. Furthermore, we model only additive microbial effects (host phenotype are not affected by for instance interactions among microbes, or microbial diversity). Fitness *R* of phenotype *i* follows a Gaussian distribution and is calculated as:

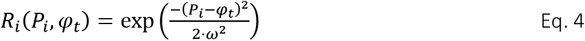

Here, *ω*^2^ is the selection strength toward the optimum phenotype, which is defined by the time-specific environment *φ*_*t*_ (see above). The optimum phenotype thus varies through time, and 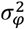 determines how much it varies (Eq. 2). Note that the average environment is set at 0 (Eq. 2), which matches the average microbial effect (Eq. 1) and thus the average host phenotype (Eq. 3). This implies that hosts with the average microbiome, are well-adapted on the long-term. We thus purposely did not include any directional selection on the mean phenotype. We sample *N* hosts (with replacement) with probabilities scaled to their fitness, to select hosts that reproduce. This procedure assumes non-overlapping host generations.

Next, microbes of reproducing hosts are vertically transmitted. Parameter *τ* (ranging between 0-1) controls vertical transmission fidelity, and is the core parameter that we vary in order to assess its effects on long-term host fitness. For each microbe that is present in a reproducing host, we sample whether or not it is transmitted to the offspring based on transmission probability *τ* (blue box in Fig. S1A). We then complement the community with randomly sampled microbes from the source pool, to keep the total number of microbes within each host equal to 100. Larger *τ* values thus indicate more faithful transmission; *τ* = 1 results in strict vertical transmission, where offspring is born with the same microbiome composition as its parent upon reproducing. In contrast, *τ* = 0 corresponds to a scenario where offspring is born with a microbiome that is fully sampled from the environmental source pool. We define this process as ‘horizontal transmission’. We note that the microbial composition in the environmental source pool does not change over time, differing from the model developed by Roughgarden (2020), where also the environmental source pool responds to selection. After this, microbial dynamics within the next host generation take place as explained above.

### Microbiome transmission fidelity shapes host phenotypic distributions and fitness

In order to evaluate how microbiome transmission fidelity (*τ*) affects phenotype distributions and fitness in a host population (Fig. S1B), here and below, we ran simulations for 1000 host generations while following 500 hosts. Including the last 500 host generations, ensuring that these numbers result in robust results, we then calculated the average phenotypic squared mean 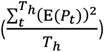, variance 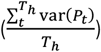 and fitness 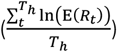. We start by setting *T*_*m*_ at 1 (extended below) to consider the simple scenario in which microbial composition does not change during host development.

More faithful vertical microbe transmission decreases host phenotypic variance (Fig. 3A): when all microbes are faithfully transmitted from parents to offspring (*τ* =1), all phenotypic variance disappears due to a combination of selective (loss of hosts that are mal-adapted in that time-step) and stochastic (loss of hosts by chance) events. This is analogous to a population in which, in the absence of new mutations, all genetic variation eventually disappears (Appendix S2). On the other hand, when *τ*=0, each host starts with completely random set of microbial species every host generation, resulting in maximal phenotypic variance (analogous to a –biologically unrealistic-scenario where each allele mutates every generation; Appendix S2).

**Figure 3:**
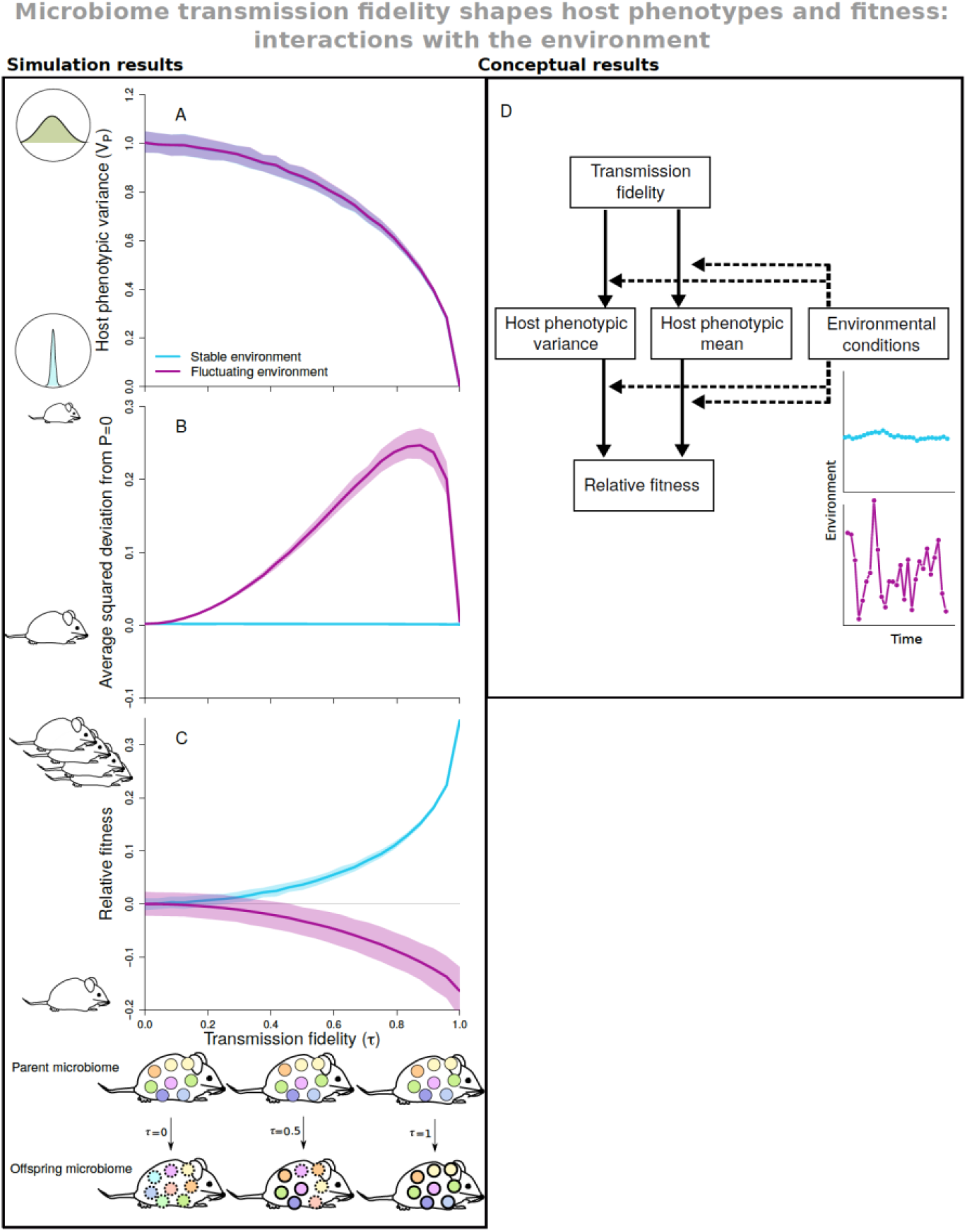
Vertical microbiome transmission fidelity affects both host phenotypic variance (A), and the deviation from the optimal mean (B), which together shape long-term fitness (C). The environment shapes these relations (blue: stable environment, purple: fluctuating environment; note that lines completely overlap in A). D) Conceptual overview of these results. Microbiome composition does not change during host development, and phenotypes are solely determined by a host’s microbiome. We ran simulations for 1000 host generations, and results in A-C are based on the last 500 host generations. In B), the squared deviation is calculated by comparing host phenotypes to the long-term optimum phenotype of 0. In C), fitness is calculated by comparing host phenotypes to the time-specific environment using Eq. 4. Lines show median values based on 250 replicate simulations. Shaded regions indicate 68% ranges of the simulations. Illustrations next to the y-axes illustrate: A) the amount of phenotypic variation among hosts, ranging from no variation to high variation; B) the average population-level deviation from the optimal phenotype (with arbitrary body sizes); C) relative fitness (compared to long-term fitness when *τ* =0), ranging from low (here depicted as one host offspring) to high (here depicted as many offspring). Transmission fidelity (x-axis) is defined as the expected proportion of microbes that is faithfully transmitted from parents to offspring, here illustrated for *τ* =0, *τ* =0.5 and *τ* =1. *ω*^2^ = 1; 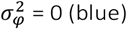 or 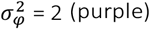 ; *V*_*α*_ = 0.01

A non-zero transmission fidelity not only reduces the amount of variation, but also has the potential to shift the population-level mean phenotype from one generation to the next. This, under constant selection, results in the average phenotype matching the long-term phenotype optimum of 0 (Fig. 3B). However, under fluctuating selection, faithful transmission increases the average deviation (per time-step) from the long-term phenotype optimum of 0 (Fig. 3B), up to when transmission fidelity is almost 100%. We note that we simulated a large host population, consisting of 500 individuals. Reducing the population size makes the population more sensitive to stochastic processes, leading to an increase in the number of mal-adapted populations that deviate from the optimum of 0 (Appendix S3).

These effects of transmission fidelity on host phenotypic variance and mean, translate into differences in long-term host fitness (Fig. 3C). As expected, in a stable environment, strict faithful transmission maximizes fitness by reducing phenotypic variance while ensuring that the average phenotype matches the optimum (blue lines in Fig. 3A-C). In contrast, under sufficiently large fluctuating selection, fitness decreases with an increasing transmission fidelity (purple line in Fig. 3C).

These effects on long-term fitness are driven by both the effects *τ* has on the amount of phenotypic variation, and on deviations from the optimal phenotype (Fig. 3D). This implies that under fluctuating selection, even when the average phenotype is kept at its long-term optimum, hosts benefit from not (or only partly) transmitting their microbiome as a means to increase phenotypic variation (see Appendix S4 for results when the mean is kept at 0). In other words, under fluctuating environments, genotypes that produce offspring with randomly assembled microbiomes, attain a higher long-term fitness compared to genotypes that faithfully transmit (at least part of) their microbiome. This is because long-term fitness, as it is a multiplicative process, is very sensitive to occasional low values: only one year with a fitness of 0, is enough to reduce long-term fitness to 0. Therefore, a strategy that reduces the variance in fitness across generations, by enhancing variation within one generation, can be adaptive in the long run, despite a reduction in an individual’s expected fitness (Philippi & Seger 1989; Bruijning *et al.* 2020) (Fig. 2). We here show that host genotypes can express such a strategy by lowering their microbiome transmission fidelity.

A mathematical framework to calculate the optimal amount of phenotypic variation as a function of the strength of stabilizing selection and how much it fluctuates through time, is proposed by Bull (1987) (Appendix S5). When keeping the average phenotype at 0, our results match Bull’s predictions (Appendix S4).

### High transmission fidelity enables adaptive tracking in predictable environments

Up to this point, we considered no temporal autocorrelation, i.e. the environment at time *t* was not predictive for the environment at time *t+1*. In nature, however, many environmental conditions are temporally autocorrelated (Halley 1996; Vasseur & Yodzis 2004), and the combination of environmental predictability and variation could shape host evolutionary responses (Botero *et al.* 2015). We increase the temporal autocorrelation (predictability *ρ*) up to 0.999, adding directional selection on the mean phenotype. We assess the optimal transmission fidelity for combinations of environmental variance *σ*^2^ and predictability *ρ*, keeping the average environment at 0 (Fig. 4). Optimal transmission fidelity is assessed by comparing long-term fitness of populations that differ in their transmission fidelity. When there is considerable environmental variation and a low environmental predictability (labeled with 2 in Fig. 4, see also Fig. 3), selection favors a low transmission fidelity, as this ensures i) that phenotypic variation across hosts is maintained, and ii) that the mean phenotype remains at its long-term optimum of 0. In contrast, a highly predictable environment (with the same environmental variance) (labeled 3 in Fig. 4), favors a transmission fidelity close to 1, as this allows hosts to follow changes in the mean environment through adaptive tracking. Note that, even though the environment changes in a highly predictable way, strict faithful transmission (*τ* = 1) again quickly results in the loss of all variation, hampering the population to track these changes, and strongly reduces fitness.

**Figure 4:**
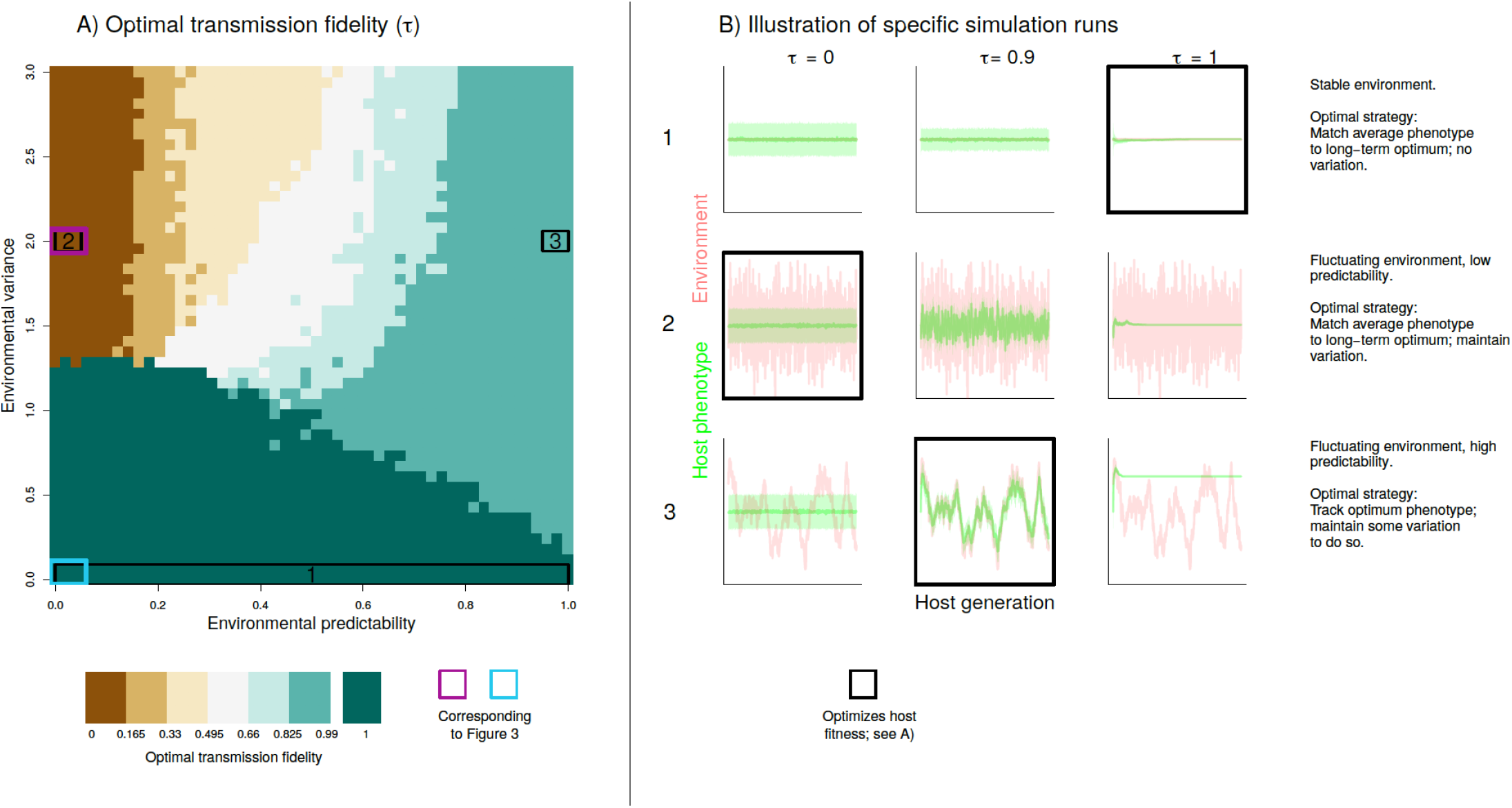
Environmental predictability and variance together shape the optimal transmission fidelity. A) Optimal transmission fidelity for 50*×*50 combinations of environmental variance and predictability (i.e. temporal autocorrelation). Colored rectangles correspond to the scenarios depicted in Fig. 3 (blue: stable environment, purple: fluctuating environment). Rectangles with labels 1 -3 indicate combinations that are further illustrated in B). These graphs show the output of specific simulation runs. Green lines indicate population-level average phenotypes, and shadings indicate one standard deviation below and above the average. Note that when the environment is stable (scenario 1), the flat red line is not visible in the graphs. Each set of graphs shows results when transmission fidelity *τ* is set at 0, 0.9 and 1, where black rectangles indicate the transmission fidelity that maximizes fitness. *ω*^2^ = 1; *V*_*α*_ = 0.01.

### Other sources of phenotypic variation among hosts

Until now, phenotypic variation among hosts was solely determined by variation in their microbiome composition, i.e. we focused only on determinants and consequences of phenotypic variance associated with the microbiome. However, phenotypic variation in natural populations is evidently not only caused by variation in microbiome composition. We will now explore what happens if we include other sources of phenotypic variation. According to a quantitative genetic framework, phenotypes can be described as the sum of one or more genetic and non-genetic components. We recently proposed an approach to framing how a microbiome contribution can be incorporated (Henry *et al.* 2019). Assuming no interactions, host phenotypes can be written as:

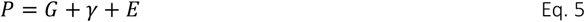

*G* is a host’s genetic value (assuming only additive effects), *γ* is its microbiome contribution (here calculated according to Eq. 3, which also assumes only additive effects), and *E* is a residual component. Total phenotypic host variance can now be written as:

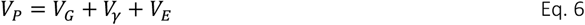

Up to this point, we have assumed that all *G* and *E* values are zero, resulting in *V*_*G*_ = *V*_*E*_ = 0, and hence: *V*_*P*_ = *V*_*γ*_. Thus, the microbiome variance required in order to optimize phenotypic variance is: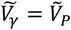. Eq. 6 illustrates that a non-zero contribution of other variance components affects the optimal microbiome variance, as now: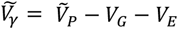. The optimal microbiome variance thus decreases with larger contributions of other sources of variation. Interestingly, lowering *V*_*γ*_ can be achieved by increasing vertical transmission fidelity *τ* (see Fig. 3a). Indeed, varying *V*_*E*_ changes the selection on transmission fidelity: larger values of *V*_*E*_ increase the optimal transmission fidelity, while weakening selection on *τ* in general (Fig. 5).

**Figure 5:**
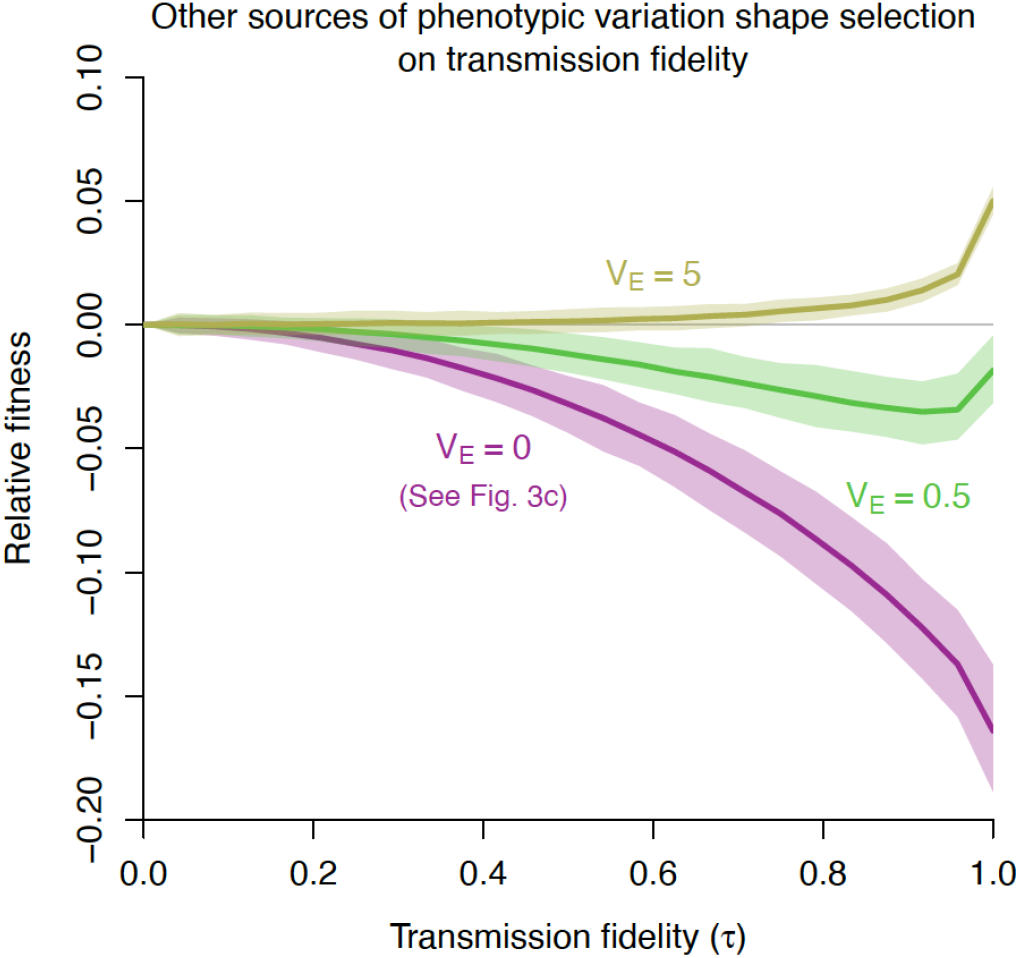
Including other sources of phenotypic variation among hosts changes selection on transmission fidelity. Colors show different values for the environmental (residual) contribution (*V*_*E*_) to host phenotypic variance. Larger *V*_*E*_ values weaken selection on vertical transmission and select for more faithful microbe transmission. In accordance with Fig. 3, we set *ω*^2^ = 1; 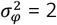 *V*_*α*_ = 0.01. Lines show median predictions based on 250 simulations; shaded regions indicate 68% ranges of the simulations

Here, we consider one focal host genotype (or strategy), such that *V*_*G*_ = 0. By the same logic as above, increasing *V*_*G*_ (due to e.g. sexual reproduction) will decrease the optimal microbiome variance, therefore increasing the optimal transmission fidelity. These findings illustrate that selection on microbiome transmission fidelity is shaped by how inherently stochastic host phenotypes are. Hosts with phenotypes with a strong stochastic component might, even under fluctuating selection, benefit from strict microbiome control, to avoid increasing phenotypic variance even more. In contrast, host with phenotypes that show little inherent variation (e.g. in the absence of environmental or genetic variation), might, under sufficiently large fluctuating selection, benefit from extra variance induced by noisy transmission.

### Changes in microbiome composition during host development

As the generation time of microbes is generally orders of magnitude shorter than that of their host, microbiome composition generally changes over the course of a host’s life (Burns *et al.* 2016; Kolodny *et al.* 2019). To account for this, we will allow neutral dynamics to affect microbiome composition between the moment hosts are born and the moment they reproduce. We do so by varying the number of microbial generations *T*_*m*_, as a measure for the relative host generation time, and varying the balance between the acquiring of new microbes from the environment (with probability *c*) and within-host proliferation (with probability 1-*c*). Parameter *c* is a measure for how much microbiome composition changes due to horizontal transmission during host development, and empirical estimates of *c* vary within the full 0 to 1 range (Sieber *et al.* 2019).

More microbial generations within one host generation (i.e. higher *T*_*m*_ values) increase phenotypic variation among hosts (Fig. 6A-C). Increasing colonization from the environment reduces the effects of *T*_*m*_ and *τ* and creates a more homogeneous phenotypic variance landscape (Fig. 6C). Thus, environmental colonization *c* has the ability to both increase and decrease phenotypic variance (Fig. 6D), depending on the transmission fidelity and the number of microbial generations occurring within a host (colored dots in Fig. 6A-C mapping onto curves on panel Fig. 6D). Both increasing the number of microbial generations and increasing colonization can strongly reduce microbiome heritability, even under strict vertical transmission, illustrating the difference between inheritance and heritability (Appendix S6).

**Figure 6:**
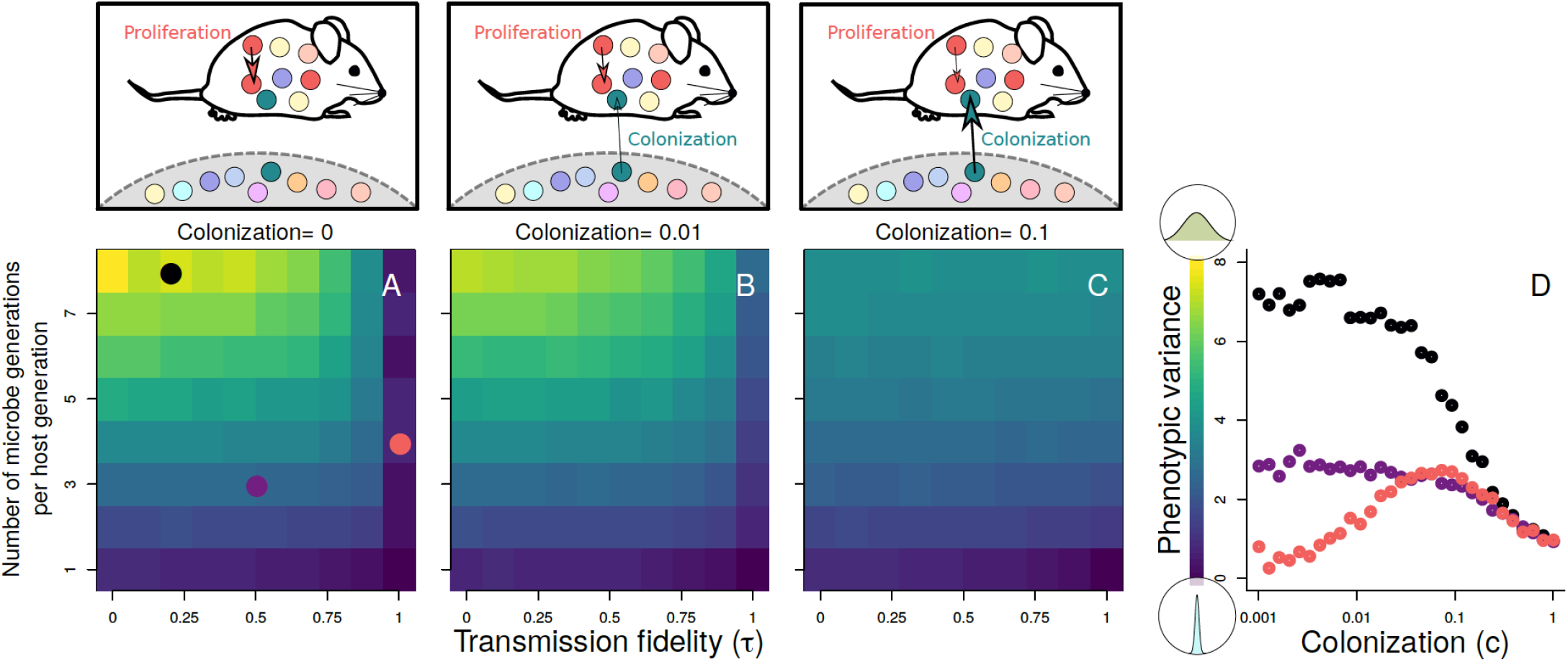
Environmental colonization and within-host proliferation alter host phenotypic variance. A-C) Effects of changesin microbiome composition during host development on host phenotypic variance, for different colonization probabilities. Dots in A) depict combinations of *T*_*m*_ and *τ* that are further explored in D). D) Phenotypic variance as a function of colonization probabilities (note the log scale on the x-axis). *ω*^2^ = 1; 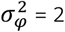 *V*_*α*_ = 0.01

These neutral dynamics *within* one host generation result in patterns as expected from ecological metacommunity theory: increased microbe (i.e. species) migration from the source pool to individual hosts (i.e. communities), decreases variation among hosts (i.e. β-diversity) (Appendix S7). This is in line with empirical studies showing that dispersal among hosts leads to more similar microbial communities (Burns *et al.* 2017; Moeller *et al.* 2018). However, as our focus is on the long-term evolutionary consequences for a population of hosts, we additionally model stochastic and selective host reproduction. This has no analog in a metacommunity framework, where communities do not duplicate or disappear. Our extended framework, including these host dynamics, shows that, even though environmental transmission generally reduces variation *within* one host generation (Appendix S7; Fig. 6D), it is required to maintain variation on host evolutionary time scales (Fig. 3A).

## Discussion

Given the importance of the microbiome for host fitness, hosts are expected to be under selection to control their microbiome composition. Here, we focused on the host feature of controlling the transfer of microbes from parents to offspring. The wide diversity in microbiome transmission fidelity that exists in nature (Fig. 1), suggests that selection tunes vertical transmission fidelity differently across host systems, yet a conceptual framework to lay out expectations for selective pressures on microbiome transmission has been lacking.

Here, we developed a general model to evaluate how vertical transmission fidelity could affect long-term host fitness, accounting for the important but often neglected feature of fluctuating environmental conditions. We found that transmission fidelity affects the amount of microbiome variation across hosts: strong control over transmission reduces variation in microbial composition across hosts, while weak control increases microbial variation (Fig. 3). We find that there are conditions under which lower transmission fidelity is beneficial, and showed that both external properties, such as the strength of phenotypic selection (Appendix S2), how selection fluctuates through time (Figs 3,4) and the importance of microbial transmission from the environment (Fig. 6), as well as host properties such as relative generation time (Fig. 6) and the relative contribution of microbiome variation to host phenotypic variation (Fig. 5), can all shape selection on microbiome transmission fidelity. Our results suggest that, under unpredictable environmental conditions, imperfect transmission can be adaptive, not only by affecting the mean host phenotype (Fig. 3B), but also by tuning phenotypic variability among hosts (Fig. 3A,D). In this study, we focused on selection on *vertical* transmission fidelity, and defined *horizontal* transmission as the process of picking up random microbes from the environment. However, we note that heritable control over horizontal transmission (Picazo *et al.* 2019) (with variation among host genotypes in how faithfully they pick up the same microbes as their parent), results in essentially the same population-level outcomes as heritable control over vertical transmission. We thus expect selection on heritable horizontal transmission to be consistent with the presented results here.

### Environmentally-dependent microbial effects

Our model provides some general predictions on selection on control over microbiome transmission. Importantly, we illustrate that environmentally-dependent microbial effects are crucial for selection to favour intermediate transmission fidelity: selection in the case of consistently beneficial microbes always favours fully faithful transmission (whereas selection favours no transmission for consistently detrimental microbes). Indeed, such environmental-dependent microbial effects on hosts are found in a range of systems, suggesting that mixed modes of inheritance might indeed be favoured in nature. For example, in thrips, the effects of *Erwinia* bacteria depend on its host diet (De Vries *et al.* 2004), proposed as an explanation for why thrips did not evolve strict vertical transmission (De Vries *et al.* 2004). Mycorrhizal effects in plants can depend on environmental conditions (Johnson *et al.* 1997), and in damselfish, the benefits of cleaning gobies depend on the presence of ectoparasites (Cheney & Côté 2005). Finally, in aphids, fitness effects of multiple symbionts (and their environmental interaction), as well as transmission patterns, are relatively well understood (Russell & Moran 2006; Oliver *et al.* 2010). For instance, the maternally transmitted facultative symbiont *Hamiltonella defensa* provides protection against endoparasitoid wasps (Oliver *et al.* 2003), but comes with an apparent fitness cost in parasitism-free environments (Oliver *et al.* 2008). Indeed, variation in selection can maintain hosts both with and without this symbiont (Ives *et al.* 2020). *Serratia symbiotica*, another facultative aphid symbiont, increases host heat tolerance (Chen *et al.* 2000; Montllor *et al.* 2002; Russell & Moran 2006), but can decrease host fitness under lower temperatures (Chen *et al.* 2000).

### Understanding the lack of faithful transmission of beneficial microbes

Despite the numerous examples of microbes with environmentally-dependent effects, there are also microbes with seemingly consistent benefits for their hosts, irrespective of the environment, but lacking faithful transmission. For instance, *Burkholderia* bacteria provide insecticide resistance to their host bean bugs (Kikuchi *et al.* 2012) and we are not aware of any study reporting negative host effects of this symbiont. While, based on our model, we predict that, in such cases, host-level selection should favour perfect vertical transmission, *Burkholderia* bacteria are not vertically transmitted, and hosts have to reacquire their symbiont from the soil every generation (Kikuchi & Yumoto 2013). We have the following three explanations for the lack of faithful transmission of consistently beneficial microbes.

First, developmental or physiological constraints can make it difficult to ensure full concordance between parent and offspring microbiome composition. It might be difficult to faithfully transmit all microbial species from parents to offspring, and furthermore, microbial dynamics often have ample of opportunity to change microbiome composition during host development (Burns *et al.* 2016; Kolodny *et al.* 2019), due to stochastic processes or competition among microbes. This is especially true for hosts with longer generation times (Fig. 6).

Second, there could be hidden fitness costs of the microbe, i.e. decreasing host fitness components in certain life stages or in certain environments. For instance, *Drosophila* individuals from which their microbiome has been removed, suffer from a reduced fecundity (implying beneficial microbial effects); however, their life span is increased (Gould *et al.* 2018). Whether or not we expect selection to favour sterile or non-sterile flies in this case, depends on the sensitivity of fitness to changes in fecundity and life span (Caswell 1978), i.e. on the life history of the species. Although this particular selective outcome (sterile versus unsterile) is unlikely to be relevant in nature, it illustrates that the microbiome can have opposing effects on different fitness components, which could explain why apparent beneficial microbes are not always faithfully transmitted. In our model, we considered only one fitness component. Future studies that combine our model with demographic models, such as matrix population models or Integral Projection Models (Caswell 2001; Ellner & Rees 2006), could help us better understand selection patterns on microbiome transmission in species with more complex demography.

Third, weak selection on the optimal vertical transmission fidelity could explain the lack of faithful transmission. This could be due to different reasons. In humans, mothers and newborn babies share a similar microbiome, however, this similarity breaks down over the first few years (Asnicar *et al.* 2017; Ferretti *et al.* 2018; Yassour *et al.* 2018). This may partly reflect constraints due to the long host development time (see first explanation), but it may additionally be that these maternally transmitted microbes are particularly important early in life, with the importance disappearing in later life stages, weakening selection on parent-offspring microbiome resemblance later in life. Selection on transmission also becomes weaker with a lower contribution of microbial variation to host phenotypic variation (Fig. 5). Furthermore, host fitness could be relatively insensitive to the phenotype shaped by the microbiome, weakening the strength of selection on microbe transmission. We modelled host fitness as a function of a single phenotypic trait, while in reality, fitness results from the combined effects of many phenotypic traits. Again, future studies combining our approach with more elaborated demographic models could further explore this. Finally, the broader demographic context of a species can affect selection on the optimal amount of variation in one phenotypic trait (Bruijning *et al.* 2020).

### Understanding faithful transmission of detrimental microbes

Similarly puzzling is why hosts would faithfully transmit microbes that seem consistently disadvantageous (or neutral) for host fitness. For instance, *Wolbachia* infections are very common in insects, and *Wolbachia* is transmitted through strict vertical maternal transmission. By manipulating host reproduction, many *Wolbachia* groups are considered to be parasites (Werren *et al.* 2008). Why would hosts transmit such harmful microbes? Using the same reasoning as above, there could be hidden benefits of the microbe, benefiting certain life stages or fitness components, or under certain environmental conditions. For example, in flies, *Wolbachia* can block the establishment of viral pathogens (Teixeira *et al.* 2008), where a higher *Wolbachia* density leads to better protection (Chrostek *et al.* 2013). However, in the absence of viral pressure, high densities lead to earlier death in flies (Chrostek & Teixeira 2015). This again illustrates the potential complexity in environmental-dependent fitness effects, affecting multiple components of fitness.

An alternative explanation is selection at the level of the microbe. Especially if host-level selection on transmission fidelity is low (see above), microbe-level selection might efficiently increase transmission rates up to a certain extent. A recent modelling study illustrates this idea, although their focus was not on transmission fidelity: weaker host-level selection increased the success of faster-growing neutral microbes (van Vliet & Doebeli 2019). Expanding our model to include different microbe strategies, instead of the neutral microbial dynamics that we included, could provide novel insights in how strong microbe-versus host-level selection must be for non-zero transmission rates of pathogenic microbes to evolve.

### Going forward

Developing approaches that balance realistic complexity with tractable simplicity is a crucial challenge in advancing our theoretical understanding of microbiome-host dynamics. The conceptual similarity between microbiome transmission fidelity and genetic mutations outlined here, resulted in a close match between our simulation results and quantitative genetic predictions (Appendix S2; see also Ravel et al., 1997). This illustrates the potential power of leveraging the wealth of theory existing in the field of quantitative genetics. For instance, could we use developed theory on epistatic variance (e.g. Hansen, 2013) to inform us on how host-microbe interactions affect selection on transmission? What are the consequences of such interactions for coevolution of host and microbes? Evaluating where the conceptual similarity between the transmission of microbes and genetic material breaks down, and where we thus need to develop new theory, will be crucial.

In addition to the development of theory, there is a clear need for empirical data in order to test model-generated predictions on selection on vertical transmission fidelity. Despite the number of studies on transmission fidelity in different systems (Fig. 1), studies that combine such data, with information on how microbiome composition affects host fitness, in an environmentally-dependent manner, are very limited. This is not surprising; as many eukaryotic hosts have highly diverse microbiomes, it is challenging to characterize phenotypic and fitness effects of specific microbial species, let alone their interaction with the environment. Moreover, biological processes such as interactions between microbes (Gould *et al.* 2018), interactions between hosts and microbes (Liu *et al.* 2019; Oyserman *et al.* 2019), microbe- or environment-dependent transmission rates (Osaka *et al.* 2008; Gundel *et al.* 2009; Rock *et al.* 2018; Liu *et al.* 2019), and microbe-level selection (Burns *et al.* 2017; Moeller *et al.* 2018), might complicate it even further to detect host-level selection patterns on transmission fidelity in natural systems. Our model simplifies many important biological features. This simplicity is warranted as it yields insight into the core processes at play; and added biological realism is unlikely to alter our main conclusion: in unpredictable environments, host selection can favour unfaithful microbe transmission in order to increase variation among offspring, leading to a bet hedging strategy. Moreover, we show that even the limited range of biological processes that we explore here are sufficient to result in optimal transmission fidelities encompassing the wide range observed in nature. Our simple model thereby provides a starting point for generating testable predictions. With empirical data in hand to motivate such refinement, it will be straightforward to extending the model to include, e.g. non-neutral microbial dynamics.

To test our predictions, carefully designed experiments could be used. First and foremost, this requires a host system where microbiome variation translates into variation in a fitness-related host phenotype (or in a measure of fitness directly). This could be done with either entire microbiome communities (Fig. 7A) or with specific host-associated microbes (Fig. 7B). Second, there must be some environmental interaction with this phenotypic trait or with fitness, so that by varying environmental conditions, fluctuating selection can be imposed. Alternatively, artificial selection could be used, for example alternately selecting for small versus large host body sizes. Third, one must be able to control microbiome transmission fidelity from parents to offspring, and possibly the importance of microbial colonization from the environment. When these criteria are met, this will allow testing of how vertical microbial transmission fidelity affects long-term host fitness, under different selection regimes (Fig. 7).

**Figure 7:**
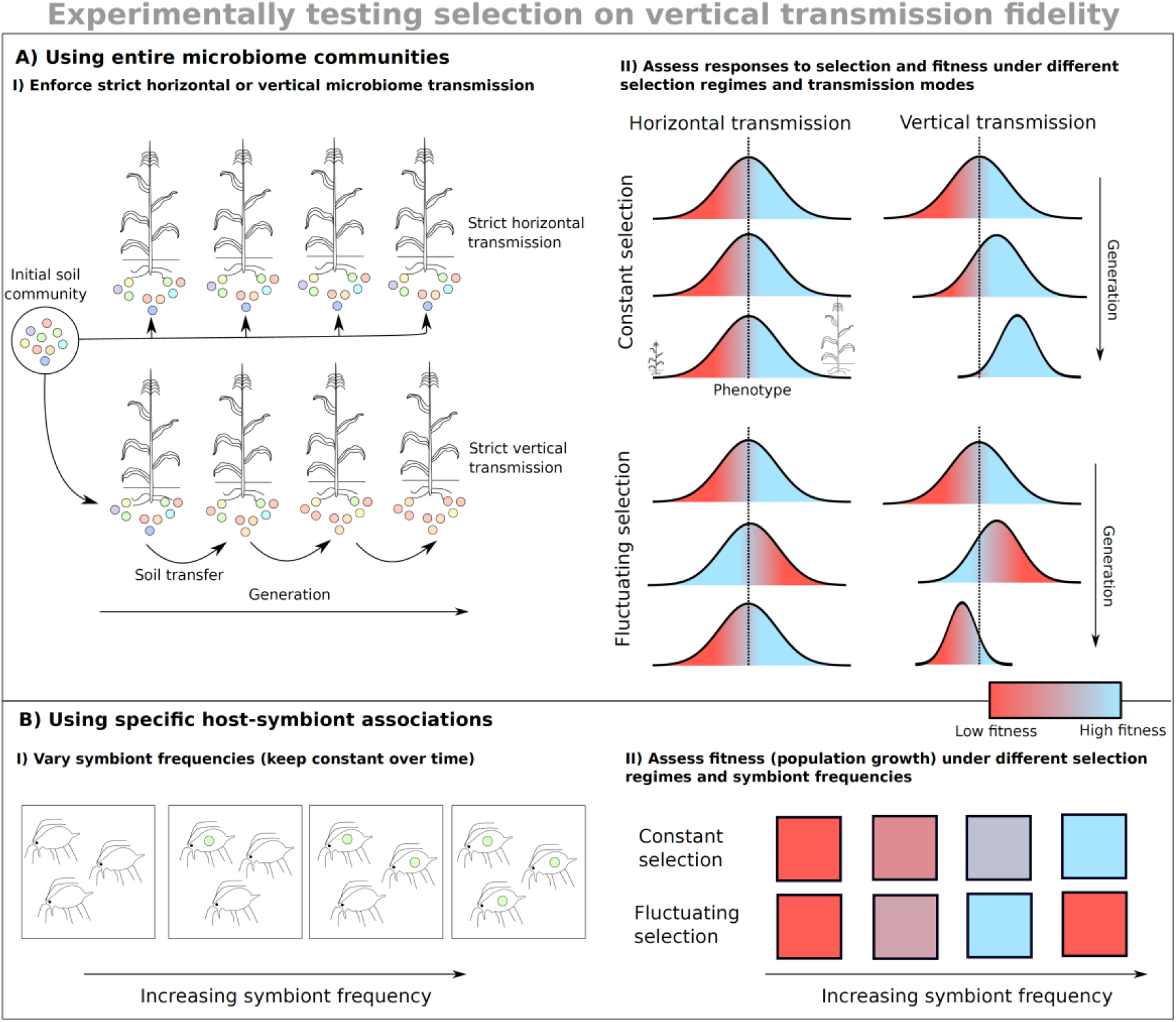
Empirical approaches to testing how microbiome transmission can affect host fitness. A) Soil transfers in plant microbiomes can be used to enforce strict environmental acquisition or vertical transmission. By either successively inoculating plant generations with their initial starting microbial community (upper row in panel A-i), or passaging the microbial community from the previous to the next generation (bottom row in panel A-i) (Morella et al. 2020), transmission of microbes can be controlled. Each host generation, artificial selection can be used to select plants based on their phenotype (e.g. plant size, illustrated here), whereby selection regimes vary (imposing either constant or fluctuating selection). Based on our results, we expect that under constant selection, strict vertical transmission increases fitness compared to strict horizontal transmission, as it allows phenotype distributions to respond to selection. In contrast, under sufficiently large fluctuating selection, vertical transmission reduces phenotypic variation, decreasing long-term fitness. B) A single microbe with a clear effect on host performance can also be used to study selection on transmission fidelity. As discussed in the manuscript, aphid fitness effects of several vertically transmitted symbionts, as well as their environmental-dependence, are quite well understood (Oliver et al. 2010). This makes aphids arguably a suitable system to study selection on vertical transmission fidelity. To do so, one could vary the symbiont frequency in different aphid populations (panel B-i). Populations can be followed through time, while keeping symbiont frequencies constant. Based on our results, we expect that under constant selection, a symbiont frequency of 100% (or 0%) optimizes population growth (panel B-ii), which can be realized by perfect vertical transmission. Under fluctuating selection, some intermediate symbiont frequency might be favored (panel B-ii), which can be achieved by noisy vertical transmission.

The low microbiome heritability in many host systems has led to a lively and ongoing discussion on the importance of the microbiome for host evolution (Moran & Sloan 2015; Douglas & Werren 2016; Theis *et al.* 2016; Henry *et al.* 2019). We propose that a low microbiome heritability resulting from imperfect transmission may actually benefit hosts under certain conditions. These conditions include an environmentally-dependent microbial effects, where effects change over time. We lay out that this is because microbiome transmission fidelity shapes phenotypic distributions in a population of hosts. The phenotypic mean and variance optimizing fitness depends on various host and environmental properties. Using a simple model, we have generated some general predictions on how faithful microbiome transmission should be, given these properties. We believe that the way forward is to test these predictions under controlled conditions, in combination with continuing the development of theory. This will provide new insights in the wide diversity in transmission modes that we observe in nature, contributing to our understanding in the role of the microbiome for host evolution.

## Acknowledgements

MB is supported by NWO Rubicon (019.192EN.017); JFA is supported by National Institutes of Health (NIH) grants GM124881; CJEM is supported by NSF DEB grant 1753993; LPH is supported by NSF-GRFP grant DGE1656466.

