## Appendix for "When the microbiome defines the host phenotype: selection on vertical transmission in varying environments"

August 27, 2020

#### Contents

**S1 Schematig overview of our model** **2**

**S2 Mutation-selection-drift models** **3**

**S3 Increased stochasticity in small populations** **5**

**S4 Changes in mean phenotype** **6**

**S5 Bull's model** **7**

**S6 Heritability vs inheritance** **9**

**S7 Microbial dynamics within one host generation** **10**

### S1 Schematig overview of our model

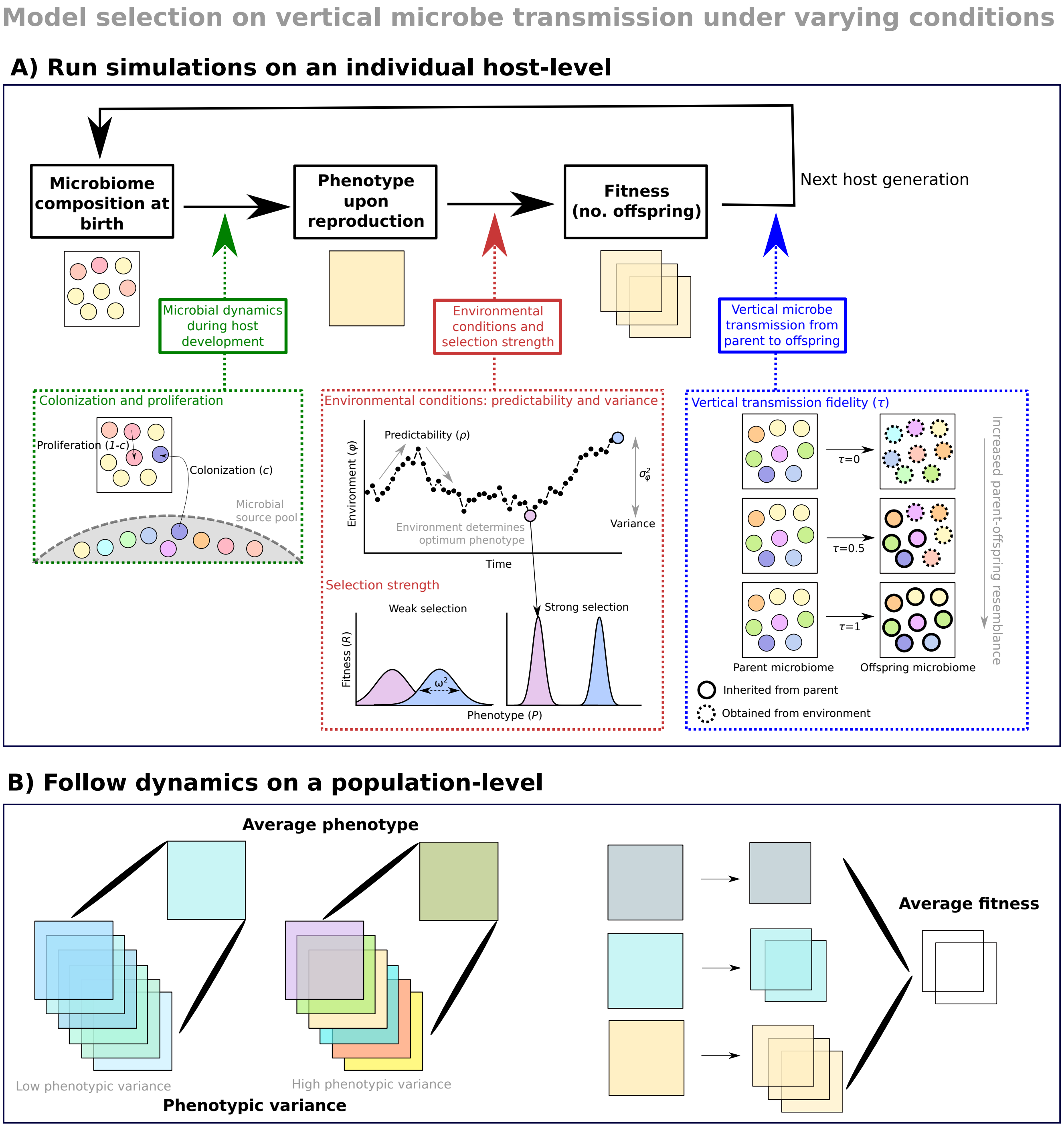


Figure S1: Schematic overview of our model. Upper panel shows the different steps of our simulations, and the parameters that we vary. Bottom panel shows the output that we obtain from each simulation.

#### S2 Mutation-selection-drift models

Mapping our models onto existing quantitative genetic models gives us a better quantitative understanding in how vertical transmission fidelity shapes variation among hosts. We here provide and briefly discuss the relevant quantitative genetic equations that we use for comparing to our model; for more background and details on mutation-selection-drift models we refer the reader to e.g. Walsh and Lynch (2018).

We start by considering a neutral trait, i.e. in the absence of selection, and assume that this trait is determined by many loci with only additive effects (i.e. no epistasis and dominance). Drift reduces the amount of genetic variation each generation, as a function of the effective population size $N_e$. New mutations, on the other hand, increase the amount of variation. The variance produced each generation due to mutations (the 'mutational variance') equals in diploid populations:

$$V_m = 2n\mu\sigma_\alpha^2 \tag{S1}$$

(Lynch and Walsh, 1998). Here, $n$ is the number of loci underlying the phenotypic trait, $\mu$ is the mutation rate, and $\sigma_\alpha^2$ is the variance of the effects of new mutations (we assume that $\mu$ and $\sigma_\alpha^2$ are constant over all loci). In our simulations, we can view each of the 100 microbes present in a host as a different allele, affecting host phenotypes in an additive way. Considering a diploid population, this is analogous to $n = 50$ loci. Microbes get faithfully transmitted with one minus the 'mutation' probability: $\tau = 1 - \mu$. With probability $\mu$, a microbe gets replaced by a microbe sampled from the environmental source pool, where microbial effects are distributed as $N(0, V_\alpha)$ (see Eq. 1 in the main text). Thus, $V_\alpha$ describes the variance of the effects of new mutations, i.e. $\sigma_\alpha^2$. Filling in Eq. S1, the variance introduced each generation due to microbiome changes is thus: $100(1-\tau)V_\alpha$.

In our simulations, newly arriving microbes ('mutants') have effects centered on 0, independent of the parental microbe. This corresponds to a House-of-Cards (HOC) mutational model (Kingman, 1978; Walsh and Lynch, 2018; Zeng and Cockerham, 1993), where each mutation has a new effect drawn from a normal distribution, and is thus independent of the prior allele state. The equilibrium additive genetic variance under the HOC model is:

$$\tilde{\sigma}_\alpha^2(HOC) = \frac{4N_eV_m}{1 + 4N_e\mu} \tag{S2}$$

(Walsh and Lynch, 2018; Zeng and Cockerham, 1993). Running our simulations without any phenotypic selection, and comparing the observed phenotypic variances to predicted equilibrium genetic variances using Eq. S2, we obtain very similar results (Fig. S2a). Variances range from 0 when $\mu = 0$ (or $\tau = 1$), to $2n\sigma_\alpha^2$ when $\mu = 1$ and $N_e \to \infty$.

The presence of stabilizing selection reduces the additive genetic variance. Fleming's prediction states that in a diploid asexual population with infinite size, the equilibrium genetic variance under the balance between mutations and stabilizing selection, is approximately:

$$\tilde{\sigma}_\alpha^2(F) = \sqrt{V_mV_s}[1 - \frac{\sqrt{V_m}}{\sqrt{V_s}}(\frac{\eta+3}{16U} - \frac{19}{16})] \tag{S3}$$

Under the assumption that $\frac{\sigma_\alpha^2}{\mu V_s} \ll 1$ (Bürger, 1998, 1999; Fleming, 1979; Turelli, 1984). $V_s$ is the strength of selection on genotypic values: $V_s = \omega^2 + V_E$. As in our simulations all phenotypic variance is due to microbiome variance, $V_s = \omega^2$. Parameter $\eta$ denotes the kurtosis of the mutation distribution and is thus 0 for normally distributed mutations. Finally, $U$ is the mutation rate of the genome: $U = 2n\mu$. A stochastic version of Eq. S3, including drift, can be written as:

$$\tilde{\sigma}_\alpha^2(SF) = \frac{-V_s}{4N_e} + \sqrt{(\frac{V_s}{4N_e})^2 + \tilde{\sigma}_\alpha^4(F)} \quad \text{(S4)}$$

(Bürger, 1999). We used Eq. S4 to calculate equilibrium genetic variances for varying selection strengths ($\omega^2$) (under a stable environment), and compared these predictions to our simulations. Again we found a very close match between prediction and observations (Fig. S2b). Fleming's approximation assumes a Random-Walk mutational model, while, as explained above, our simulations correspond to a HOC mutational model. Under a Random-Walk model, mutational effects are centered on the parental allele effects (Kimura, 1965; Walsh and Lynch, 2018; Zeng and Cockerham, 1993). In the absence of selection, this results in no upper bound on the genetic variance, as it continues to increase with the effective population size (Walsh and Lynch, 2018). Strong stabilizing selection, however, imposes an upper bound to the mutational effects (microbes with strong effects are selected against), resulting in a close match between Fleming's prediction and our simulation results, despite the difference in underlying mutational model. When selection decreases (higher $\omega^2$ values), however, Fleming's prediction overestimates our observations, while observations eventually approach the HOC mutation-drift balance predictions (Eq. S2) (Fig. S2c).

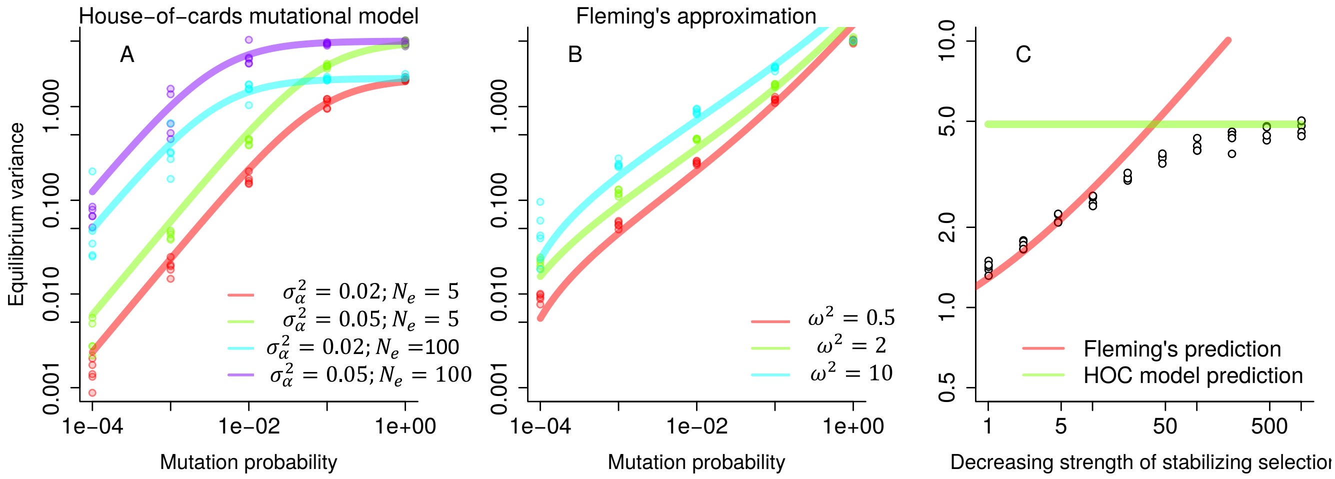


Figure S2: Comparison of our host-microbiome simulations (dots) to quantitative genetic mutation-selection-drift models (lines). A) In the absence of selection, our simulations closely resemble a House-of-Cards mutational model. B) When including stabilizing selection, Fleming's approximation adequately describes our simulations. However, as the underlying mutational model in Flemings approximation differs from a HOC mutational model, bias increases when selection becomes weaker, with our simulations approaching the HOC model in the absence of selection (C).

### S3 Increased stochasticity in small populations

In Figure 3 of the manuscript, we show how the microbiome transmission fidelity shapes host phenotype distributions. In doing so, we simulated large populations, consisting of 500 individuals, in order to obtain robust results. This results in a limited role of stochasticity, explaining the relatively low variation across replicated simulations (see shaded regions in Fig. 3A,B). In smaller populations, however, populations are, unsurprisingly, more sensitive to stochastic processes. This is illustrated in the figure below. Here, we set transmission fidelity $\tau$ at 1, implying strict vertical transmission, and assessed the average deviation from P=0 for varying population sizes. In small populations, there is an increase in the number of maladapted populations (i.e. a larger deviation from the optimal phenotype).

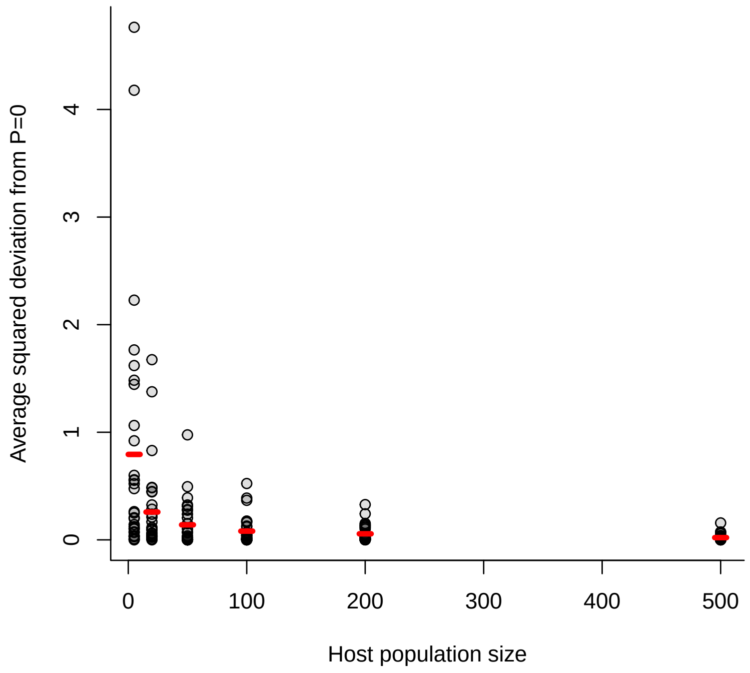


Figure S3: Relationship between host population size and the average squared deviation from P=0. Grey dots indicate individual simulations (30 per population size), grey lines indicate median values for each population size. $\tau$=1; $\omega^2 = 1$; $\sigma^2_\varphi = 2$; $V_\alpha = 0.01$.

#### S4 Changes in mean phenotype

In Figure 3 of the manuscript, we show how the microbiome transmission fidelity shapes host phenotype distributions, which translate into effects on long-term host fitness. We show that under constant environmental conditititions, faithful transmission optimizes host fitness. In contrast, under fluctuating conditions, low vertical transmission results in the highest long-term fitness. We here show that these effects remain, even if we keep the mean phenotype constant (at 0) for both environmental conditions. This was done by manually mean-centering phenotypes in each time step, by subtracting each phenotype by the average time-specific phenotype. This illustrates that, under fluctuating selection, host benefit from not (or only partly) transmitting their microbiome, as a means to increase phenotypic variation.

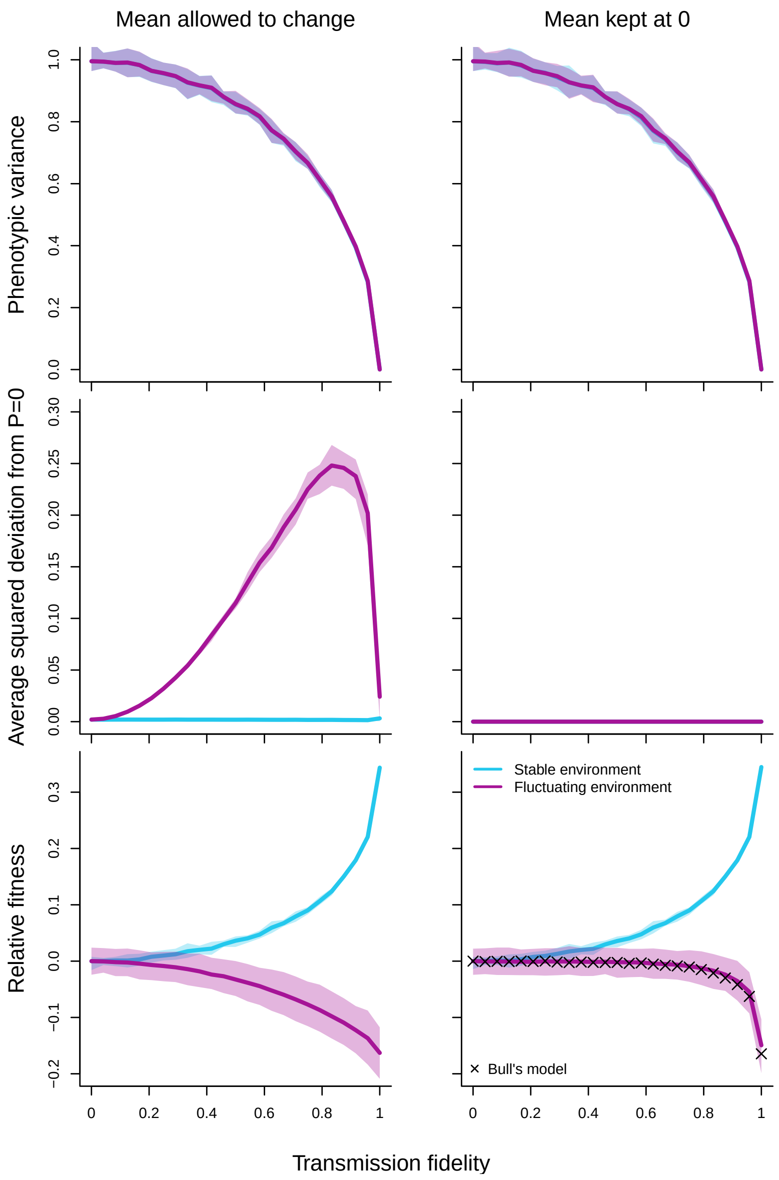


Figure S4: Relationship between transmission fidelity and phenotypic variance (upper row), deviation from the long-term optimal mean (middle row) and long-term fitness (bottom row) when selection can shift the mean phenotype (left; corresponds to Fig. 3 in the manuscript) or when keeping the mean phenotype fixed at 0 (right). When we keep the mean host phenotype at 0, we can use Bull's modeling framework (Bull, 1987) to calculate long-term fitness (crosses in bottom right panel), based on the relation between transmission fidelity and phenotypic variance.

### S5 Bull's model

Bull (1987) presents a mathematical model to calculate the optimal phenotypic variance ($\tilde{V_P}$), for the same fitness function as we evaluate in our paper, showing that $\tilde{V_P}$ is a function of both the environmental variance $\sigma^2_\varphi$ and the stabilizing selection strength $\omega^2$:

$$\tilde{V_p} = \begin{cases} 0 & \text{if } \sigma_\varphi^2 < \omega^2 \\ \sigma_\varphi^2 - \omega^2 & \text{otherwise} \end{cases} \tag{S5}$$

(Bull, 1987). This holds as long as the average phenotype matches the optimal phenotype in the average environment ($\overline{P} = \overline{\varphi}$). In our simulations, the average phenotype across all time steps is indeed 0 (Fig. S4). This is expected, as both the average microbial effect and the long-term optimal phenotype equal 0. However, when $\tau > 0$, selection has the potential to push the mean away from 0 from one host generation to the next (see Fig. 3b of the manuscript). For example, if at one time point, selection favors larger phenotypes, hosts that possess microbes with more positive effects will have a higher chance of reproducing. As part of the microbiome is faithfully transmitted, more of these microbes with positive effects will be transmitted, altering the mean phenotype in the next generation. Indeed, although the average phenotype across all time steps is zero, the squared deviation from zero within time steps is larger at higher transmission fidelities, up to a certain point (Fig. 3b of the manuscript). This implies that faithful microbiome transmission comes with a fitness cost under fluctuating environments, not only by decreasing phenotypic variance, but potentially also by short-term selection pushing the average phenotype away from the optimum. When we do keep the mean equal to 0, the optimal transmission fidelity resulting in the highest long-term fitness, matches the transmission fidelity generating the optimal phenotypic variance using Bull's model prediction (Eq. S5) (Fig. S4).

#### S6 Heritability vs inheritance

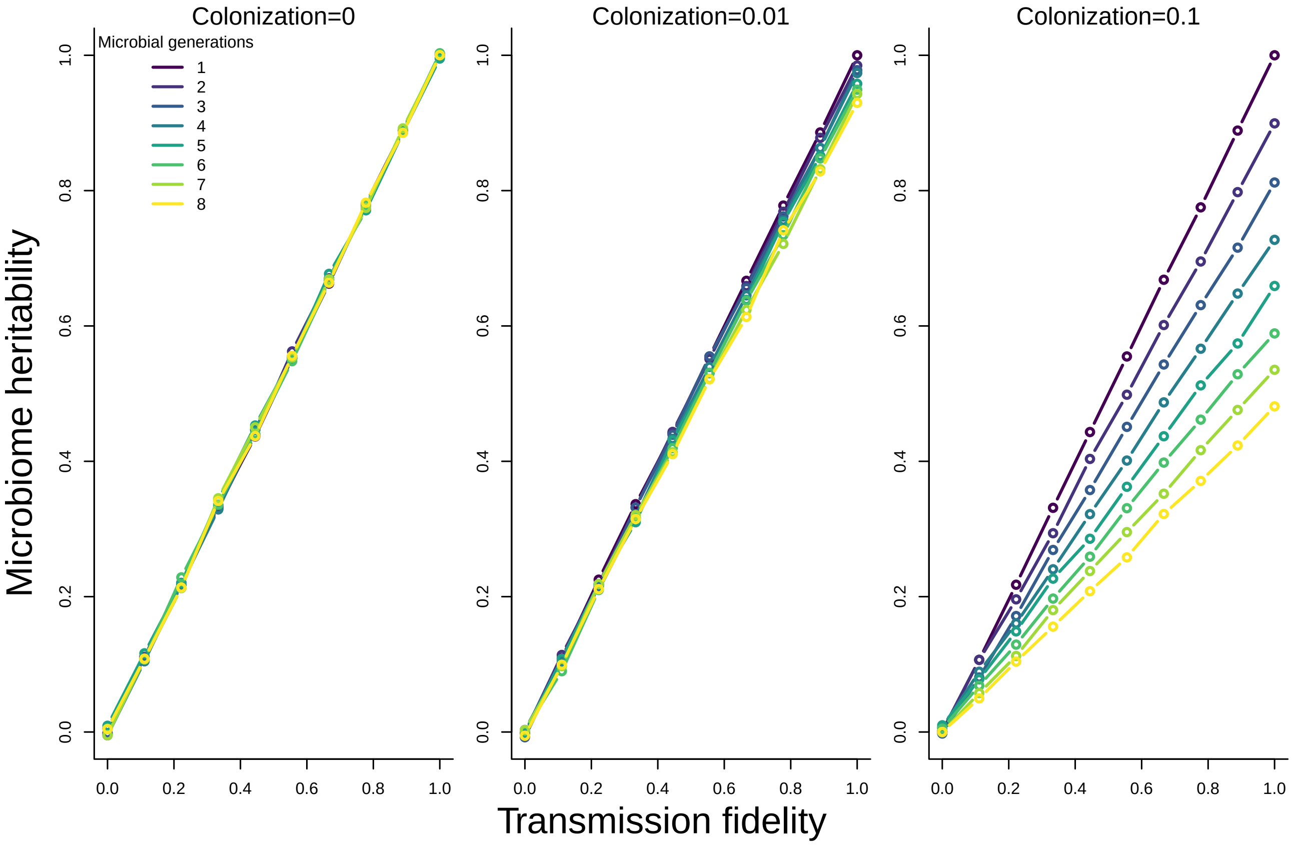


Figure S5: Heritability (averaged across 5 replicated simulations) is a function of the transmission fidelity, colonization from the environment, and the numberof microbial generations within a host generation. Heritability is measured as the slope of a regression between parent and offspring phenotypes upon the moment of reproduction, averaged across time steps. If there is only one microbial generation within a host generation and/or without colonization, the heritability equals the transmission fidelity. However, when one or both increase, heritability decreases, illustrating the difference between inheritance and heritability.

### S7 Microbial dynamics within one host generation

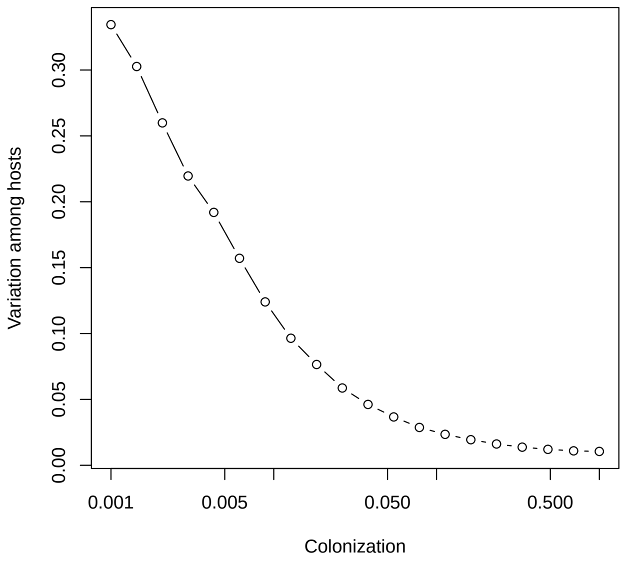


Figure S6: Within one host generation, increased colonization from the microbial source pool decreases microbial variation among hosts, as predicted from metacommunity theory. Variation among hosts is calculated as the average microbial diversity within each host, divided by the total microbial diversity across all hosts.
